# MetaQuad: Shared Informative Variants Discovery in Metagenomic Samples

**DOI:** 10.1101/2023.10.09.561628

**Authors:** Sheng Xu, Daniel C. Morgan, Gordon Qian, Yuanhua Huang, Joshua W. K. Ho

**Affiliations:** School of Biomedical Sciences, Li Ka Shing Faculty of Medicine, The University of Hong Kong, Pokfulam, Hong Kong SAR, China; Laboratory of Data Discovery for Health Limited (D^2^4H), Hong Kong Science Park, Hong Kong SAR, China

## Abstract

**Motivation:** Strain-level analysis of metagenomic data has garnered significant interest in recent years. Microbial single nucleotide polymorphisms (SNPs) are genomic variants that can reflect strain-level differences within a microbial species. The diversity and emergence of SNPs in microbial genomes may reveal evolutionary history and environmental adaptation in microbial populations. However, efficient discovery of shared polymorphic variants in a large collection metagenomic samples remains a computational challenge.

**Results:** MetaQuad employs a density-based clustering technique to efficiently differentiate shared variants from non-polymorphic sites using shotgun metagenomic data. Empirical comparisons with other state-of-the-art methods show that MetaQuad significantly reduces the number of false-positive SNPs without greatly affecting the true-positive rate. We used MetaQuad to identify antibiotic-associated variants in patients who underwent *Helicobacter pylori* eradication therapy. MetaQuad detected 7,591 variants across 529 antibiotic resistance genes. The nucleotide diversity of some genes is increased six weeks after antibiotic treatment, potentially indicating the role of these genes in specific antibiotic treatments.

**Availability:** MetaQuad is an open-source Python package available via https://github.com/holab-hku/MetaQuad.

**Supplementary information:** Supplementary data are available at *XXXX* online.

## 1 Introduction

Strain-level identification is a popular modern metagenomics technique (Marx, 2016). A bacterial strain is a genetic variant or a subtype within a bacterial species. Variants are essential mechanisms for bacteria to adapt and survive, allowing for natural selection to occur. For example, in the gut microbiome community, antibiotics can have a significant impact, species may need to adapt to survive (Koo *et al*., 2019) . Understanding bacterial strains and their adaptations is a critical aspect of microbiota investigation (Yan *et al*., 2020). Microbes can exhibit an array of phenotypic properties with minor changes in their DNA code. Contradiction may arise in some microbiome literature, when different labs detect opposing associations with certain diseases (Marx, 2016). Strain-level analysis of the microbiome composition may provide an additional layer of information that may lead to more robust and reproducible results.

Strain-level analysis is feasible with sequencing reads because a single nucleotide polymorphism (SNP) substitution reflects the difference in strain level. Shotgun metagenomic sequencing has greater power than 16S rRNA gene sequencing to detect less abundant bacterial species (Durazzi *et al*., 2021), granting it the ability to detect SNPs. However, successfully identifying strain level difference has long been a problem, especially for short-read sequencing data. Misalignment of short sequence reads may lead to false positive SNP calls. It can produce high-confidence SNPs, which are hard to filter out by variant calling algorithms, as they are likely to be supported by multiple reads together with high mapping quality (Day-Williams and Zeggini, 2011). Sequencing error furthermore confounds detection of low-frequency SNPs. Excessive false positive SNPs may mask the effect of true SNPs, which in turn may lead to issues in downstream analyses (Ma *et al*., 2019).

SNPs with consistent allele frequency changes within a population are of key interest, as they offer insights into evolution by natural selection (Wiberg *et al*., 2017). Here, we define this type of SNPs as shared informative variants. Many tools have been developed to study strain level differences, some of which are constructed especially for the shotgun metagenomic sequencing, such as metaSNV (Costea *et al*., 2017) and instrain (Olm *et al*., 2021). However, to our knowledge, no tools have been developed to jointly detect informative variants from a population of samples, due in part to the challenges of distinguishing shared informative variants from background noise. To address these limitations, we present a new computational tool called MetaQuad. MetaQuad can efficiently detect the shared informative variants, with high accuracy and precision.

## 2 Methods

### Variants calling of shotgun metagenomic data

Variant calling pipelines typically consist of three major steps. First, shotgun metagenomic sequencing reads are aligned to a database of microbial reference genomes using BWA-MEM (Li, 2013) and filtered to remove low-quality reads. Next, the aligned reads from BAM files are piled up using cellSNP-lite (v1.2.0) (Huang and Huang, 2021) to identity SNPs with minimum allele frequency (minMAF) of 0.02 within a population. The output data from cellSNP-lite usually contains a large number of microbial SNPs, including shared variants, sample-specific variants, and sequencing errors. Finally, MetaQuad selects shared informative variants using a density-based clustering technique with a minimum number of samples in each cluster. The shared informative variants can be further analyzed to calculate nucleotide diversity of genes or genomes.

### MetaQuad model

MetaQuad is a computational method to detect shared informative variants from a population of samples. MetaQuad processes allele counts of each sample output from cellsnp-lite (Huang and Huang, 2021) (Fig.1a). The pipeline prior to MetaQuad involves cellsnp-lite calculating pileup and allele counts from bulk DNA sequencing data for each variant. MetaQuad evaluates metagenomic variants using a density-based clustering technique called OPTICS (Ordering Points To Identify Cluster Structure) (Ankerst *et al*., 1999). For each variant identified by cellSNP-lite, one-dimensional OPTICS is applied based on the allele frequency (AF) of each sample. The AF for each variant in each sample is considered as a data point, and OPTICS clustering is employed to group similar data points. A minimal number of samples is required to form a cluster for each group.

**Fig. 1:**
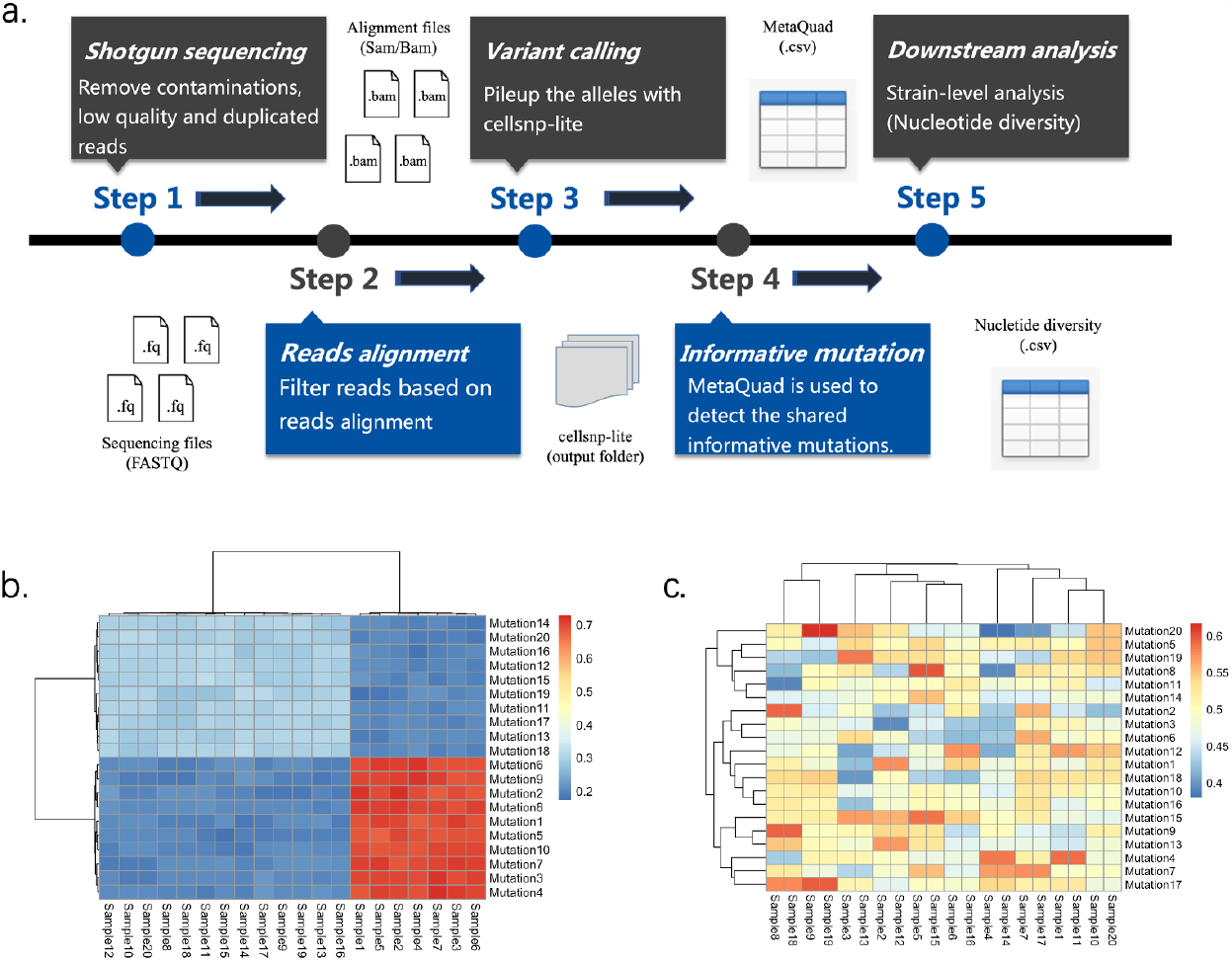
Overview of analysis pipeline and example of shared variants. **(a)** Recommended analysis pipeline for MetaQuad. The Shotgun metagenomic sequencing files are collected after removing contaminations (e.g., host contaminants), low-quality and duplicated reads. Filtered reads are aligned to a reference database using alignment tools, and low mapping quality reads are further filtered out. Cellsnp-lite is used to count the alleles of each sample, and the output data is processed by MetaQuad. All variants are listed in a CSV file with number of clusters, which distinguishes informative variants. Informative variants can be utilized to study the nucleotide diversity of each gene or genome. **(b)** Example of informative variants. Informative variants can be found in multiple samples with consistent changes in allele frequencies across populations. The allele frequencies of informative variants are similar within each population. **(c)** Example of random variants (background noise). Random variants can be found in one or more samples, but their allele frequencies do not have a consistent change, and the frequencies can vary greatly between samples. In the figure, different colors represent different allele frequencies.

The OPTICS algorithm consists of four main steps:

1. Reachability distance calculation: Compute the reachability distance between each point and its neighboring points.
2. Core distance calculation: Determine the core distance of each point with its nearest neighbor, based on the minimal number of samples.
3. Cluster extraction: Extract clusters from regions of points with high reachability distances.
4. Cluster identification: Analyze the reachability distances of the points within each cluster to identify distinct groups.

MetaQuad outputs a table containing information on all detected variants from cellSNP-lite. Importantly, it calculates the estimated number of clusters for each variant. Shared informative variants can only be confirmed if there are at least two clusters present because SNPs with only one cluster may result from limitations in the reference database or sequencing errors and are therefore removed from the study. The output table also includes additional parameters, such as read depths (DP) and allele depths (AD), which can be helpful in filtering out false positive SNPs. The examples of informative variant and random variants (background noise) are shown in Fig 1b and Fig 1c.

### Assessing the validity of the assumptions of MetaQuad

Metaquad assumes that the allele frequencies of shared informative variants are similar within a population. In order to test the assumption, we analyzed differences across individuals using a dataset from the Broad Institute-OpenBiome Microbiome Library (BIO-ML) (Poyet *et al*., 2019), under BioProject PRJNA544527. We selected 30 shotgun metagenomic samples from three healthy individuals with different time point (10 samples each, with individuals coded as ‘am ‘, ‘an ‘, and ‘ao ‘). As certain strains should be shared between samples from the same person, so too should the variants.

We processed the data by first trimming adaptors and removing phix contamination with BBMap/bbduk.sh. To remove human contamination, we used BBMap/bbmap.sh with a minimum alignment identity parameter of 0.95, and we removed duplicated reads with PRINSEQ++ (Cantu *et al*., 2019). After these steps, we aligned the filtered reads to a reference database with BWA-MEM. Specifically, we used the coding sequences of the integrated gene catalog (IGC) (Li *et al*., 2014) database from the Human Microbiome Project as our reference. We further filtered the resulting BAM files with Samtools (Danecek *et al*., 2021), removing alternative and supplemental alignments, as well as reads with mapping quality scores less than or equal to 30 to ensure accuracy.

Given the large number of genes in the reference database, we selected only the genes with genus annotations and sample occurrence frequencies larger than 0.95 according to the IGC annotation table. After this filtering step, we identified 5160 genes and used them to study shared variants both within and across individuals.

### Evaluating the model performance with simulated data

Paired-end reads with a length of 151 bp were generated using InSilicoSeq (Gourlé *et al*., 2019) to simulate shotgun metagenomic sequencing with sequencing errors (error model: NovaSeq). The simulation process involved several steps. Initially, a small gut microbiome consisting of five species from the genus Fusobacterium was established. This microbiome was used as the reference genome for the simulation. The genus Fusobacterium is known to be present in the human gut, with some members promoting the development of colorectal cancer (Bi *et al*., 2022). To generate the reference genome, five Fusobacterium genomes were downloaded from NCBI (https://www.ncbi.nlm.nih.gov) with the NCBI Reference Sequence: NZ_LN831027.1 *Fusobacterium nucleatum*, NZ_JADRGD010000001.1 *Fusobacterium necrophorum*, NZ_CAEUHP010000001.1 *Fusobacterium mortiferum*, NZ_CP028103.1 *Fusobacterium varium* and NZ_CP028105.1 *Fusobacterium ulcerans*. Four strains were generated based on the original genome, with different substitution rates (Supplementary Fig. S1a). The genome variants were performed using mutate.sh from BBmap (Bushnell, 2014).

Using the genomes, the simulation was performed on datasets consisting of 100 samples divided into three groups with different percentages of strains (Fig. 2d). The number of simulated informative variants was 7,638, and each dataset contained 100 samples divided into three groups (Groups 1, 2, and 3). The detailed simulation process is described in Supplementary Fig. S1. In order to better detect the performance of MetaQuad, multiple datasets were simulated with different mean depths of the variants. The mean depths varied from 3 to 35.5, and the detailed information of each dataset is shown in Supplementary table T1.

**Fig. 2:**
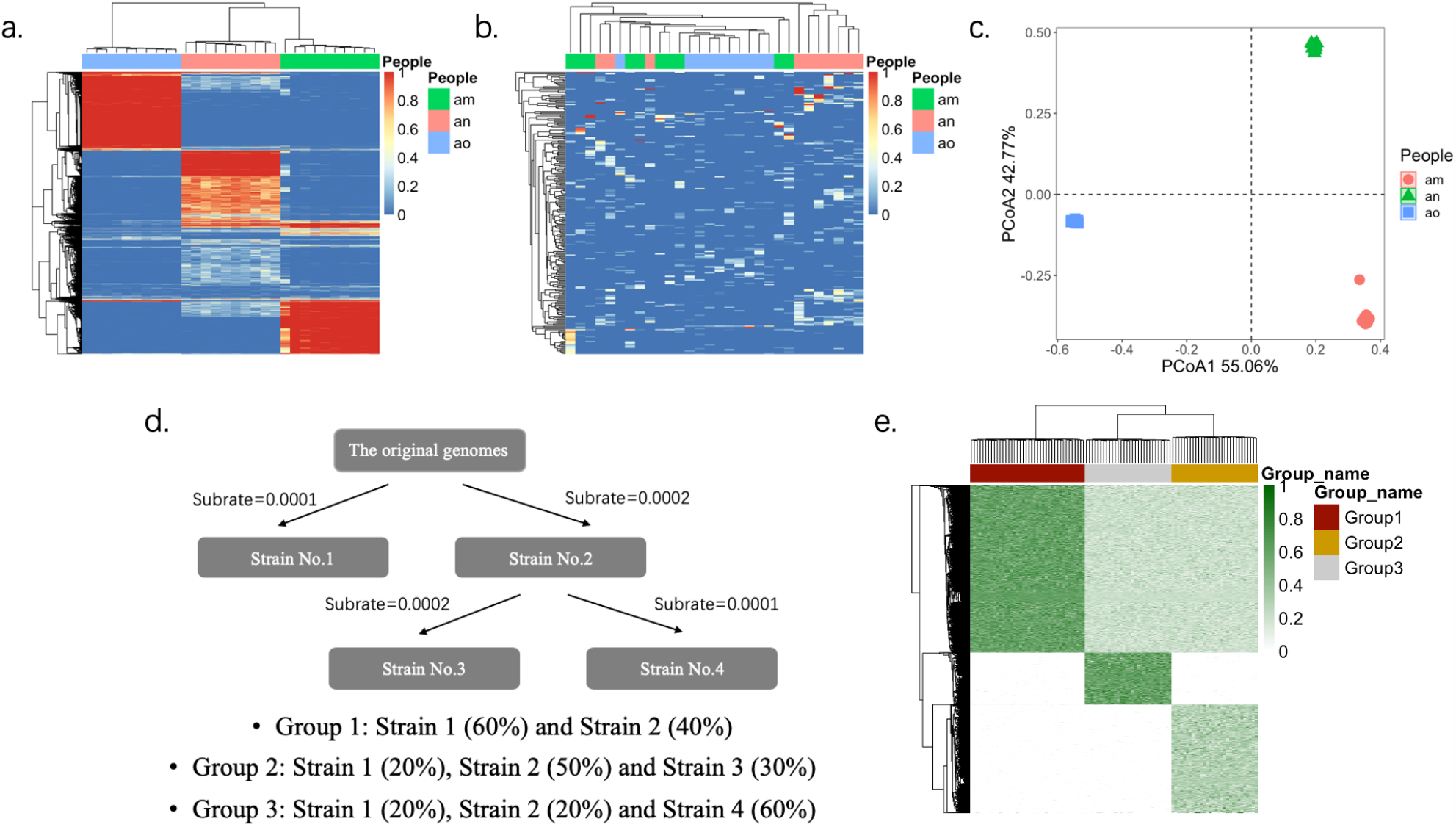
Shared informative variants in real and simulated datesets. **(a)** Allele frequencies of informative variants detected by MetaQuad within each individual. Colors indicate varying allele frequencies. **(b)** Allele frequencies of uninformative variants within each individual. Colors denote distinct allele frequencies. **(c)** Principal Coordinate Analysis (PCoA) plot of informative variants detected by MetaQuad for each individual. **(d)** Schematic representation of the strain simulation pipeline for the simulated dataset. **(e)** Allele frequencies of simulated informative variants.

The simulated reads were trimmed using BBMap/bbduk.sh to remove bases with quality scores below 10. The reads were then aligned back to the reference genome using BWA-MEM. Variant callings were performed using different tools with their default parameters, and their performances were compared later. These tools included BCFtools (Danecek *et al*., 2021), GATK’s HaplotypeCaller (Poplin *et al*., 2017), VarScan2 (Koboldt *et al*., 2012), MetaSNV (Costea *et al*., 2017) and In-Strain (Olm *et al*., 2021). The performance of these tools was evaluated using F1 score, true positive rate and false positive.

### Identification of shared variants after antibiotic treatment

Data from a study on *Helicobacter pylori* infected patients with antibiotic therapy (Wang *et al*., 2022) was used as a biological benchmark (ARG dataset). A total of 121 stool samples were collected and metagenomic shotgun sequencing was performed on these samples. The data were processed the same way as before (30 samples from three individuals).

To identify genes relevant to antibiotic resistance, we aligned the protein sequences of the IGC database to the Comprehensive Antibiotic Resistance Database (Alcock *et al*., 2019) using DIAMOND (Buchfink *et al*., 2021). We selected genes that aligned to the CARD database with an e-value less than e-50, resulting in 4948 antibiotic resistance genes (ARG genes). We then filtered the bam files to include only reads from these ARG genes, resulting in final ARG bam files that were used for further analysis, including variant calling with different methods.

We used cellsnp-lite with a minMAF of 0.02 and minCOUNT of 100, and MetaQuad with a minSample of 6, mean AD of 0.1, and mean DP of 0.5 for variant calling. All other variant calling tools were applied with their default parameters. Additional details of the data processing can be found in Figure 4a.

### Nucleotide diversity

Nucleotide diversity was calculated using allele frequencies of shared variants calculated by the output of MetaQuad and cellsnp-lite:

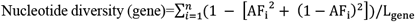

Suppose MetaQuad detected n shared variants from a gene or a genomic region. For each shared informative variant, the allele frequency (AF) could be calculated as the alternate allele (AD) divided by read depth (DP). The nucleotide diversity was further normalized by the length of genes (L_gene_). The range of nucleotide diversity was between 0 and 1, where the larger value indicated more variant accumulations.

### Statistical methods

The OPTICS algorithm was implemented using the Python scikit-learn library. Principal component analysis (PCA) was calculated using the prcomp function in R (version 4.0.2) and plotted using the ggbiplot function. Multilevel pairwise comparison was performed using the pairwise.adonis function from the ‘pairwiseAdonis’ package. The distance matrix was calculated using Bray-Curtis dissimilarity, and the P-values were adjusted using the Bonferroni method.

## 3 Results

### Consistency of shared informative variants within each individual or group

MetaQuad was initially tested on the BIO-ML dataset, which consisted of 30 samples from 3 individuals. Since the samples were collected from healthy people, the commonalities between samples from different individuals should be much smaller than those from the same individual. When analyzed with cellsnp, 67241 variants were detected, with a minimum minor allele frequency (minMAF) of 0.02. Out of all the variants, 64682 had at least two clusters with a minimum sample size of 3, while the remaining variants could only be classified into a single cluster.

For MetaQuad, variants with a minimum of two clusters are considered informative. Our analysis demonstrated that the shared informative variants were highly consistent within each individual, but not across different individuals (Fig. 2a and Fig. 2b). Variants with only one cluster were considered likely to be noise because they were not consistent within each person (Supplementary Figure 1).These results highlight that MetaQuad can effectively identify informative variants that distinguish between individuals (Fig. 3c).

**Fig. 3:**
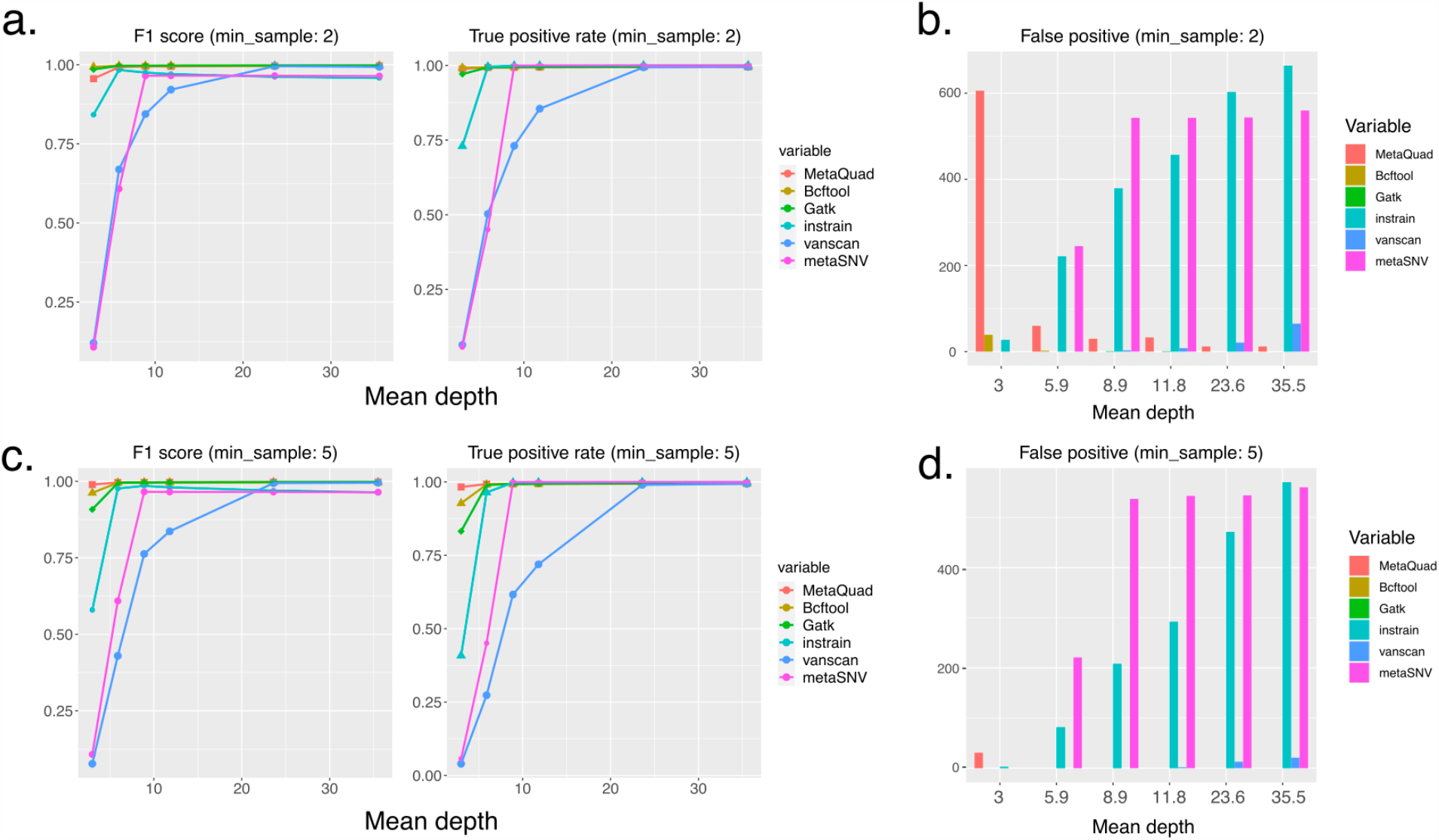
Comparison of variant calling tools. **(a)** F1 score and true positive rate of all variant calling tools, with a minimum sample threshold (min_sample) of 2. **(b)** False positive variants reported by all tools, with a minimum sample threshold (min_sample) of 2. **(c)** F1 score and true positive rate of all variant calling tools, with a minimum sample threshold (min_sample) of 5. **(d)** False positive variants reported by all tools, with a minimum sample threshold (min_sample) of 5.

### MetaQuad is a robust statistical method to identify shared informative variants from shotgun metagenomic data

MetaQuad was initially benchmarked using simulated datasets, where paired-end reads were generated from InSilicoSeq (Gourlé *et al*., 2019) (Methods). These datasets were organized into three groups, with each group containing samples that had varying percentages of strains from the original genomes (Methods, Fig. 3d). The allele frequencies of the simulated shared informative variants were highly consistent within each group, making the simulation more reflective of real-world scenarios. To thoroughly assess MetaQuad ‘s performance, multiple datasets were generated with different mean depths for the informative variants. The mean depths ranged from 3 to 35.5, and the specific details for each dataset can be found in Supplementary Table T1.

To further validate the performance of MetaQuad, we compared it with several variant calling tools (details are provided in the Methods section). We evaluated the performance of these tools using F1 scores, true positive rates, and false positives. While MetaQuad was developed to detect informative variants in a group of samples, other tools have their own applicable scenarios. To ensure a more representative comparison of the performance of different tools, we applied filters to the data. For the other tools, we considered only minor variants that appeared multiple times as informative variants, as a counterpart to MetaQuad’s minimum number of samples in the clustering. Additionally, only SNPs were considered in the comparison, and indels were removed during the filtering process. The comparative performance is shown in Fig. 3.

The combined performance of MetaQuad outperformed all the other competitors, especially with higher min-samples. All tools performed better with higher mean depths, with nearly 1 F1 scores and almost 100% true positive rates (Fig. 3a and Fig. 3c). In contrast, Varscan2 and MetaSNV could achieve high true positive rates in the case of high mean depth, but at the same time, false positive rates were also significantly increased (Fig. 3b and Fig. 3d). Bcftools and GATK performed well even with low mean depths, but they reported some sequencing errors, and they may miss some informative variants with higher min-samples and low mean depths.

Although MetaQuad may have some false positives in low mean depths, it achieved high true positive rates. However, all the false positive variants can be filtered out if we apply a filter of mean AD and DP of the data (Supplementary Fig 2). In this case, MetaQuad was better than all the other tools. If many sequencing errors and misalignments are anticipated in the study, MetaQuad will be the ideal method to apply.

### MetaQuad discovers antibiotic associated variants in human gut microbiome

MetaQuad was subsequently applied to an ARG dataset after undergoing pre-processing (methods, Fig. 4a) (Wang *et al*., 2023). The samples were divided into three groups based on their clinical treatment: CLA, LEVO, and OTHERS. Detailed clinical information can be found in the Supplementary materials (ARG_treatment_infor_modified.xlsx). The stool samples were collected from 44 patients at three distinct time points: prior to antibiotic treatment, six weeks after antibiotic treatment, and six months after antibiotic treatment. The minSample parameter of MetaQuad was set to 6, representing 5% of the total sample size. To minimize false positive variants, any variants with a mean AD less than 0.1 and DP less than 0.5 were removed.

**Fig. 4:**
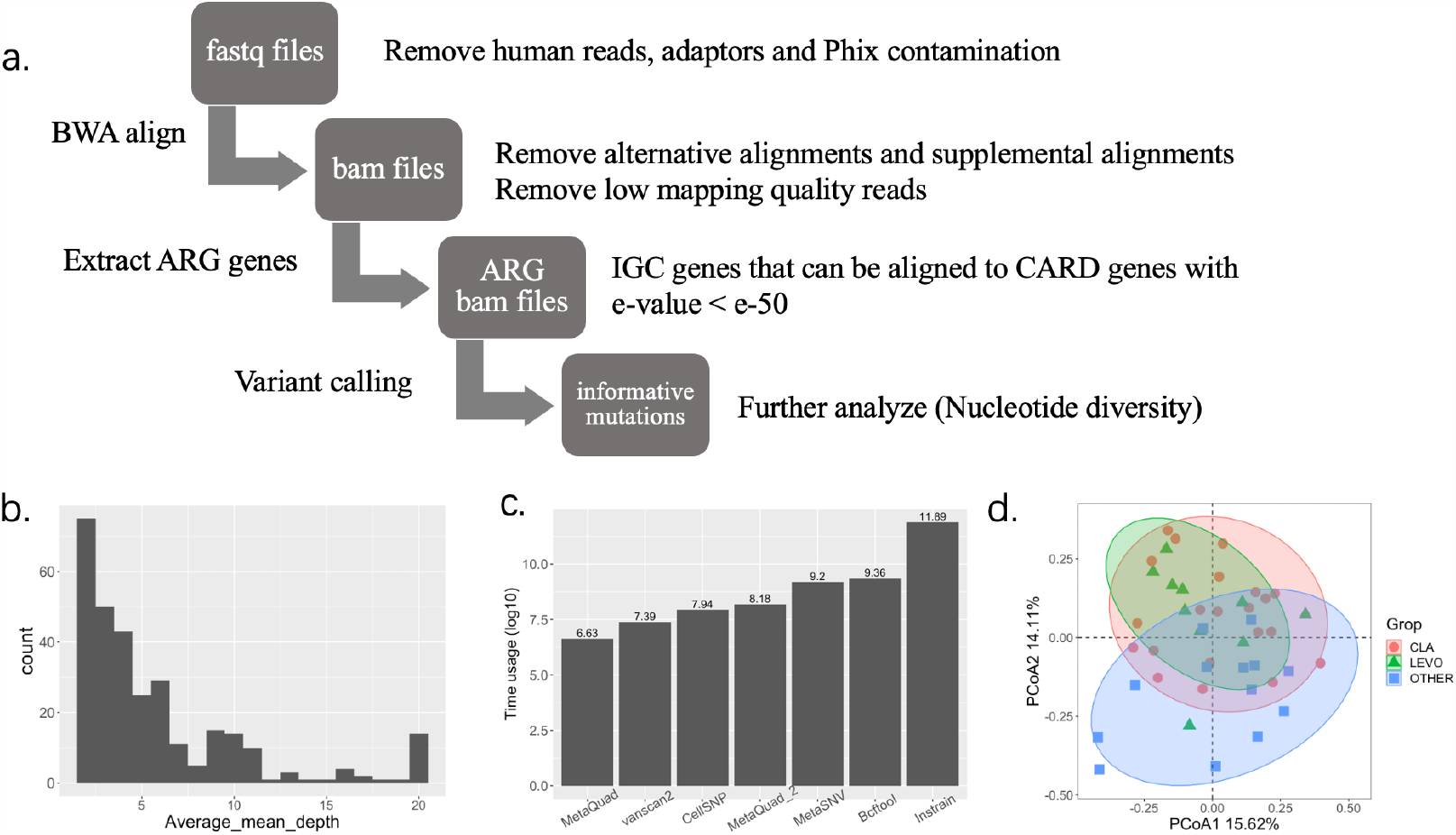
the impact of antibiotics on the human gut microbiome through antibiotic associated variants. **(a)** The pipeline used for detecting shared informative variants in the ARG dataset. **(b)** Average mean depths of each ARG gene. **(c)** the time usage of all variant calling tools in the ARG dataset, presented in log 10 transformation. MetaQuad2: total runtime of MetaQuad and cellsnp. **(d)** PcoA plot of shared informative variant six weeks after antibiotic treatment.

To investigate the impact of antibiotic treatment, we focused solely on antibiotic resistance genes (ARGs). The mean depths of the ARG genes were calculated by samtools coverage and were averaged by the total 121 samples (Fig 4b). Most ARGs had a mean depth lower than 3. Our results revealed that MetaQuad identified 7,591 informative variants from the ARG dataset, and its runtime was relatively fast, taking only about 60 minutes to identify informative variants from the ARG bam files (Fig 4c). To compare the computational efficiency of various tools, we evaluated the time usage of vancan2 and MetaSNV with cellsnp-lite and MetaQuad, as they all concurrently analyze all samples. We also considered instrain, which is specially designed for shotgun metagenomic data. For instrain, its time is calculated by its own built-in equation, and the final total time is the sum of the time required for all samples. For the other tools, we recorded the times with the unix time command (real time), using the same computational resources (ppn = 80 and storage = 500G). The MetaQuad pipeline demonstrated high efficiency, striking a balance between predictive performance and runtime costs. In fact, MetaQuad outperformed vanscan2 in our simulated dataset benchmark. Although vanscan2 was the fastest, cellsnp+MetaQuad took slightly longer, while the other tools required much longer to complete the computation.

We analyzed samples from various groups taken at different times, focusing on the allele frequencies of shared informative variants. To analyze the relationship between antibiotic treatment and the allele frequencies of these variants, we used PCoA and multilevel pairwise comparison (Fig. 4d). Prior to the analysis, we filtered out variants with NA values and calculated the allele frequencies based on the output of cellsnp-lite (AD/DP). Across all groups, we identified a total of 125, 25, and 40 shared informative variants in the samples before treatment (bT), 6 weeks after treatment (T + 6W), and six months after treatment (T + 6M), respectively. All subsequent analyses were based on these variants. Interestingly, at T + 6W, the informative variants clearly differentiated the CLA samples and LEVO samples from the others (Fig. 4d). However, this performance was not observed in the samples from the other two time points (Supplementary Fig 3). Multilevel pairwise comparison revealed that the adjusted P-values of LEVO vs. OTHER and CLA vs. OTHER were significant at T + 6W, with values of 0.03 and 0.003, respectively. However, none of these comparisons were significant at the other time points.

### Shift in Nucleotide Diversity of Antibiotic-Resistant Genes

We also examined the nucleotide diversity of antibiotic-resistant genes, which typically reflects the accumulation of variants. Informative variants were identified in 529 ARG genes, and we filtered out genes with low nucleotide diversity, leaving 15 samples with a nucleotide diversity greater than 0.005. We then focused on the remaining 13 genes to investigate changes in nucleotide diversity before and after antibiotic treatment using the Wilcoxon test. In patients treated with LEVO, the nucleotide diversity of 7 genes showed significant differences (P < 0.05) between the 00 and 01 groups (Supplementary Fig 4). The changes in nucleotide diversity were consistent across all 7 genes, with group 01 showing significantly higher nucleotide diversity than group 00, while group 00 and group 02 were very similar. These findings may help explain how bacteria adapt to antibiotic drugs, where resistance arises due to the accumulation of variants in specific resistance-determining regions (Moon *et al*., 2010). Variants accumulate under antibiotic pressure, but gradually return to pre-perturbation steady states. The new microbial genomic variants detected under antibiotic exposure are only temporary and disappear once the antibiotic pressure is removed.

## 4 Discussion

Detecting variants in shotgun metagenomic data has long been a challenging task due to the presence of common background noise. In this study, we propose and demonstrate the effectiveness of a novel analysis pipeline called MetaQuad, which utilizes shotgun metagenomic sequence samples to discover shared informative variants.

MetaQuad adopts a unique selection approach in evaluating informative variants that is more accurate than simply considering the number of samples in which variants were detected. The key strength of MetaQuad lies in its ability to detect shared informative variants even with low coverage data. This is particularly important as sequencing depth can vary across samples, making it difficult to detect variants consistently. We evaluated the performance of MetaQuad against other common variant calling tools and found that MetaQuad outperformed these tools, especially for variants with low mean depths. MetaQuad stands out from other tools with its relatively high performance and fast speed. In contrast, the other tools either perform poorly or require too much time to achieve comparable results. This makes MetaQuad a promising tool for various applications in the field.

MetaQuad is particularly relevant as researchers are increasingly focusing on strain-level differences from high-depth sequencing, making dealing with a large amount of noise inevitable. MetaQuad provides a new way to filter out useless variants and sequencing errors, thus uncovering the untapped potential of information on bacterial variants from sequencing data. Overall, MetaQuad opens up a new paradigm of analysis pipeline for shared informative variant detection in a population of shotgun metagenomic sequencing files. Its ability to detect variants even with low coverage data makes it a valuable tool for researchers working in the field of metagenomics.

## Acknowledgements

We thank Xianjie Huang for optimizing cellSNP-lite on bulk DNA-seq data, and Aaron Wing Cheung Kwok for constructive comments on adapting MQuad to MetaQuad.

## Funding

This work was supported in part by AIR@InnoHK administered by Innovation and Technology Commission of Hong Kong.

## Notes

### Competing Interest Statement

The authors have declared no competing interest.

## References

Alcock, B.P. et al. (2019) CARD 2020: antibiotic resistome surveillance with the comprehensive antibiotic resistance database. Nucleic Acids Res., gkz935.

Ankerst, M. et al. (1999) OPTICS: ordering points to identify the clustering structure. ACM SIGMOD Rec., 28, 49–60.

Bi, D. et al. (2022) Profiling Fusobacterium infection at high taxonomic resolution reveals lineage-specific correlations in colorectal cancer. Nat. Commun., 13, 3336.

Buchfink, B. et al. (2021) Sensitive protein alignments at tree-of-life scale using DIAMOND. Nat. Methods, 18, 366–368.

Bushnell, B. (2014) BBMap: A Fast, Accurate, Splice-Aware Aligner.

Cantu, V.A. et al. (2019) PRINSEQ++, a multi-threaded tool for fast and efficient quality control and preprocessing of sequencing datasets PeerJ Preprints.

Costea, P.I. et al. (2017) metaSNV: A tool for metagenomic strain level analysis. PLOS ONE, 12, e0182392.

Danecek, P. et al. (2021) Twelve years of SAMtools and BCFtools. GigaScience, 10, giab008.

Day-Williams, A.G. and Zeggini, E. (2011) The effect of next-generation sequencing technology on complex trait research: NEXT-GENERATION SEQUENCING AND COMPLEX TRAIT RESEARCH. Eur. J. Clin. Invest., 41, 561–567.

Durazzi, F. et al. (2021) Comparison between 16S rRNA and shotgun sequencing data for the taxonomic characterization of the gut microbiota. Sci. Rep., 11, 3030.

Gourlé, H. et al. (2019) Simulating Illumina metagenomic data with InSilicoSeq. Bioinformatics, 35, 521–522.

Huang, X. and Huang, Y. (2021) Cellsnp-lite: an efficient tool for genotyping single cells. Bioinformatics, 37, 4569–4571.

Koboldt, D.C. et al. (2012) VarScan 2: Somatic mutation and copy number alteration discovery in cancer by exome sequencing. Genome Res., 22, 568–576.

Koo, H. et al. (2019) Individualized recovery of gut microbial strains post antibiotics. Npj Biofilms Microbiomes, 5, 30.

Li, H. (2013) Aligning sequence reads, clone sequences and assembly contigs with BWA-MEM.

Li, J. et al. (2014) An integrated catalog of reference genes in the human gut microbiome. Nat. Biotechnol., 32, 834–841.

Ma, X. et al. (2019) Analysis of error profiles in deep next-generation sequencing data. Genome Biol., 20, 50.

Marx, V. (2016) Microbiology: the road to strain-level identification. Nat. Methods, 13, 401–404.

Moon, D.C. et al. (2010) Emergence of a new mutation and its accumulation in the topoisomerase IV gene confers high levels of resistance to fluoroquinolones in Escherichia coli isolates. Int. J. Antimicrob. Agents, 35, 76–79.

Olm, M.R. et al. (2021) inStrain profiles population microdiversity from metagenomic data and sensitively detects shared microbial strains. Nat. Biotechnol., 39, 727–736.

Poplin, R. et al. (2017) Scaling accurate genetic variant discovery to tens of thousands of samples Genomics.

Poyet, M. et al. (2019) A library of human gut bacterial isolates paired with longitudinal multiomics data enables mechanistic microbiome research. Nat. Med., 25, 1442–1452.

Wang, L. et al. (2023) Altered human gut virome in patients undergoing antibiotics therapy for Helicobacter pylori. Nat. Commun., 14, 2196.

Wang, L. et al. (2022) Dynamic changes in antibiotic resistance genes and gut microbiota after Helicobacter pylori eradication therapies. Helicobacter, 27.

Wiberg, R.A.W. et al. (2017) Identifying consistent allele frequency differences in studies of stratified populations. Methods Ecol. Evol., 8, 1899–1909.

Yan, Y. et al. (2020) Strain-level epidemiology of microbial communities and the human microbiome. Genome Med., 12, 71.

