## Supplemental figures and tables for "MetaQuad: Shared Informative Variants Discovery in Metagenomic Samples"

**Supplementary materials.**

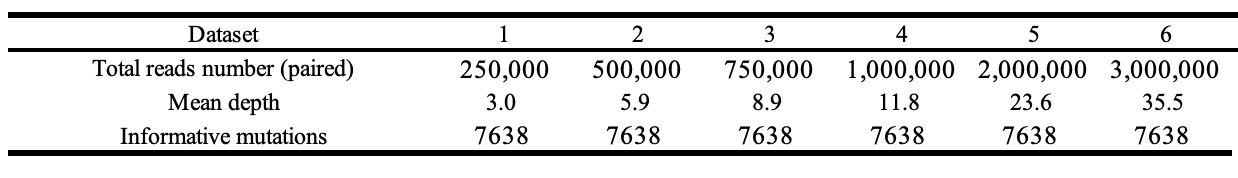

**Supplementary table T1:** Details of Simulated Datasets

Datasets were simulated with mean depth from 3.0 to 35.5, with 7628 informative mutations. Each dataset includes 100 samples with three groups.

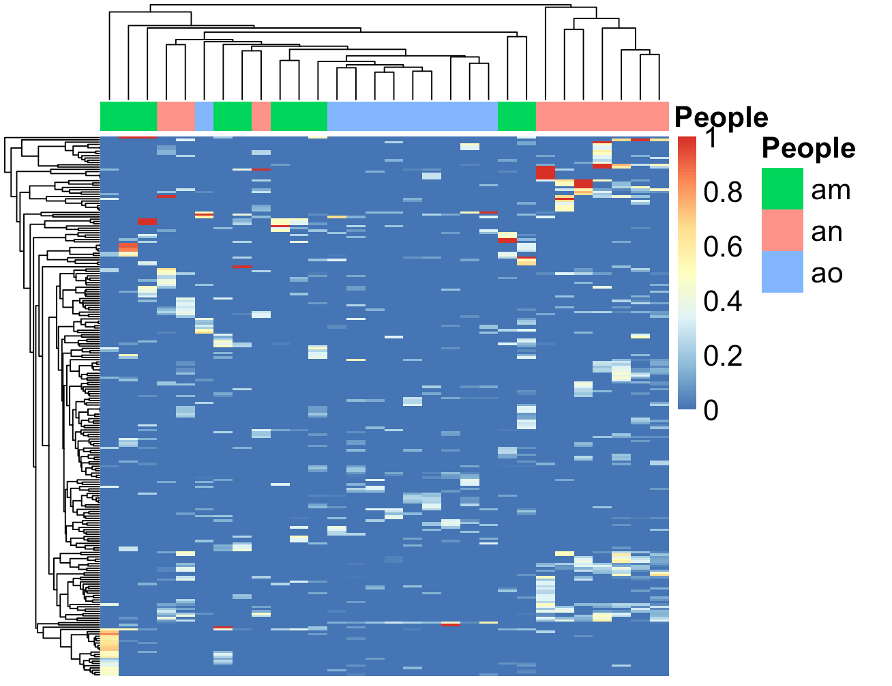

**Supplementary figure 1:** Allele frequencies of mutations with only one cluster

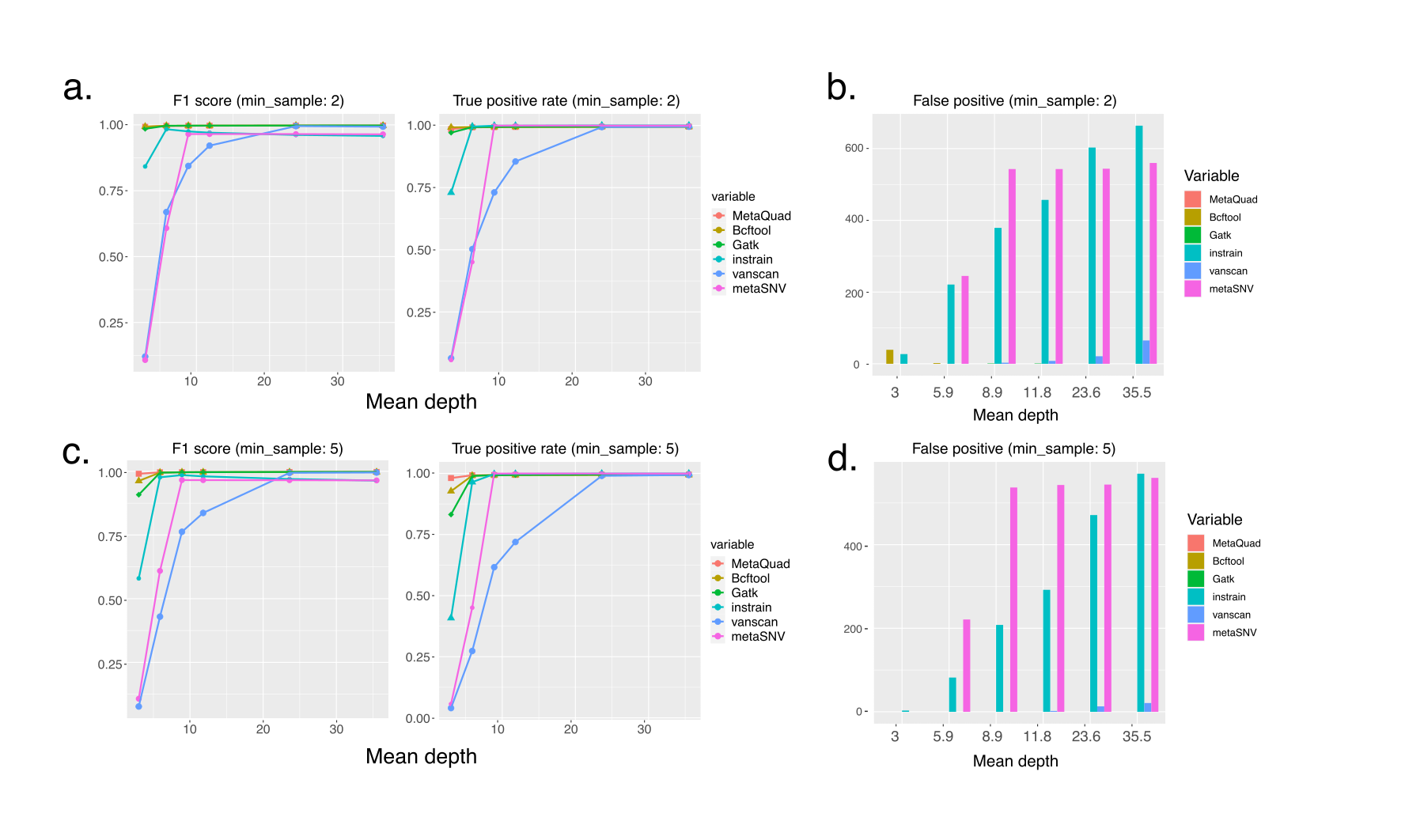

**Supplementary figure 2**: Comparison of variant calling tools with mean AD of 0.1 and DP of 0.5 filter applied for MetaQuad

(a) F1 score and true positive rate of all variant calling tools, with a minimum sample threshold (min_sample) of 2.

(b) False positive mutations reported by all tools, with a minimum sample threshold (min_sample) of 2.

(c) F1 score and true positive rate of all variant calling tools, with a minimum sample threshold (min_sample) of 5.

(d) False positive mutations reported by all tools, with a minimum sample threshold (min_sample) of 5.

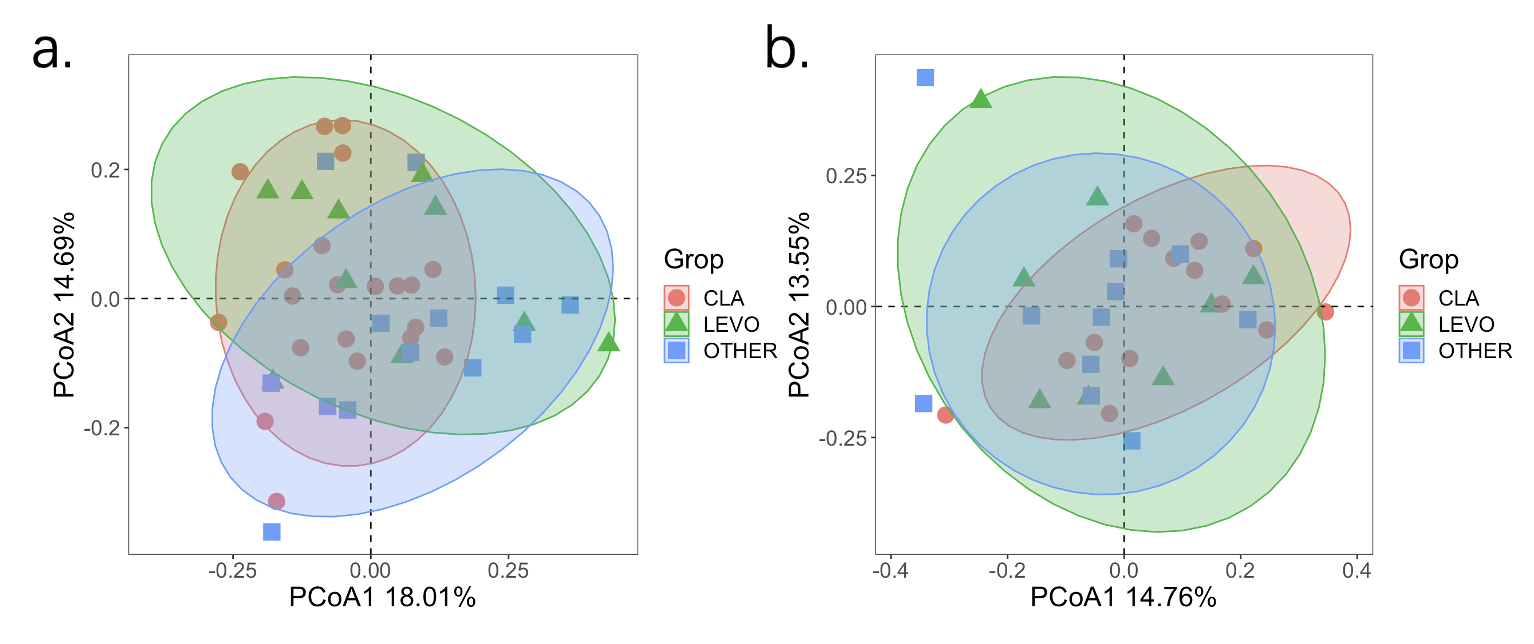

**Supplementary figure 3:** PcoA plot of shared informative mutation before (a) and six months (b) after antibiotic treatment.

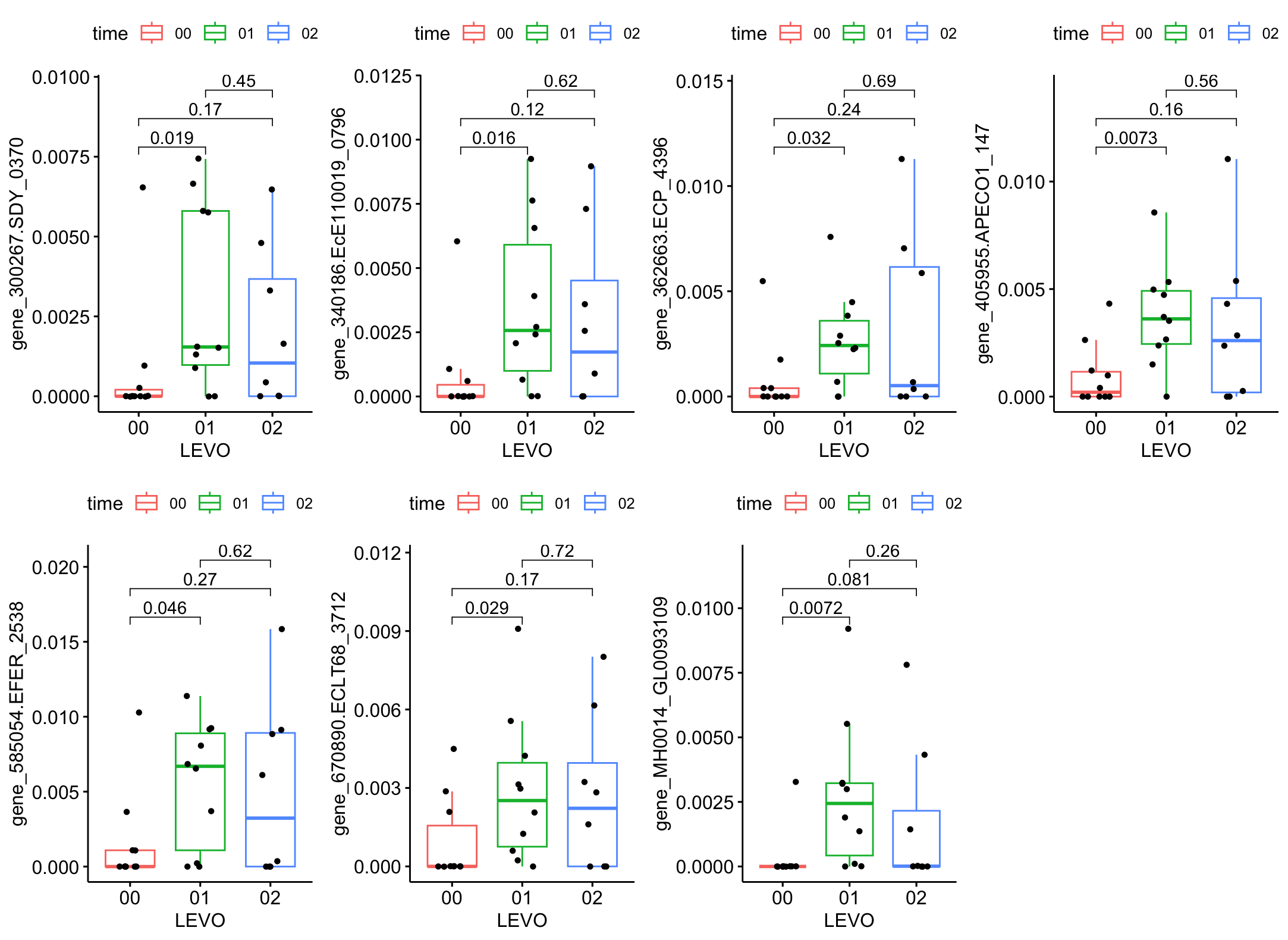

**Supplementary figure 4:** nucleotide diversity of patients with LEVO treatment.
